# Toxicity impacts on human adipose MSCs acutely exposed to Aroclor and non-Aroclor mixtures of PCBs

**DOI:** 10.1101/2022.11.17.516943

**Authors:** Riley M. Behan-Bush, Jesse N. Liszewski, Michael V. Schrodt, Bhavya Vats, Xueshu Li, Hans-Joachim Lehmler, Aloysius J. Klingelhutz, James A. Ankrum

**Author notes:** Contributed equally (Riley M. Behan-Bush, Jesse N. Liszewski).

## Abstract

PCBs accumulate in adipose where they may impact the growth and function of cells within the tissue. This is particularly concerning during adolescence when adipocytes expand rapidly. Herein we sought to understand how exposure to PCB mixtures found in U.S. schools affects human adipose mesenchymal stem/stromal cell (MSC) health and function. We investigated how exposure to Aroclor 1016 and Aroclor 1254, as well as a newly characterized non-Aroclor mixture that resembles the PCB profile found in cabinets, Cabinet Mixture, affects adipose MSC growth, viability, and function in vitro. We found that exposure to all three mixtures resulted in two distinct types of toxicity. At PCB concentrations >20 μM, the majority of MSCs die, while at 1-10 μM MSCs remained viable but display numerous alterations to their phenotype. At these sublethal concentrations, MSC rate of expansion slowed, and morphology changed. Further assessment revealed PCB-exposed MSCs had impaired adipogenesis and a modest decrease in immunosuppressive capabilities. Thus, exposure to PCB mixtures found in schools negatively impacts the health and function of adipose MSCs. This work has implications for human health due to MSCs’ role in supporting the growth and maintenance of adipose tissue.

**SYNOPSIS:** PCB mixtures found in schools are toxic to human adipose mesenchymal stem/stromal cells, stunting their growth and altering their function in ways that could contribute to metabolic diseases.

## INTRODUCTION

There is significant need to understand how exposure to polychlorinated biphenyl (PCB) mixtures found both in old and new schools contributes to the dysfunction of adipose tissue. PCBs are a group of environmental toxins containing 209 distinct congeners that were heavily produced globally from the late 1920s until being banned in 1979.^1^ Despite being banned, mixtures of different PCB congeners can still be found in capacitators (Aroclor 1016), transformers (Aroclor 1254), caulk (Aroclor 1254), and many other building materials making them ubiquitous in public spaces across the United States.^1–4^ Furthermore, evidence of Aroclor 1016 and 1254 has been found in schools as recent as 2021.^5^

While intentional production of PCBs is banned, PCBs continue to be produced as contaminants in products such as pigments and varnishes used as finishes.^6–8^ One study found finished cabinetry to be a novel, non-Aroclor source of PCB mixtures leading to elevated levels of PCBs in residential houses and apartments.^9^ It was also shown that factory workers exposed to these materials accumulate significant levels of cabinet mixture PCBs (PCB 47, PCB 51, and PCB 68) in their plasma and urine.^10^

Occupational exposure is not the only way humans are exposed to PCBs. While exposure to PCBs was once thought to be primarily through ingestion of contaminated food, it is now clear that inhalation is a major route of human exposure, specifically for the semi-volatile, lower-chlorinated PCBs.^11^ These semi-volatile PCBs do not remain fixed in place, but become volatilized over time, making them a persistent source of PCB exposure to those in the environment.^10^ Not only have non-Aroclor sources been found on small scales in residential homes, but these sources now have widespread distribution in the air of places such as Chicago despite not being manufactured in high levels prior to the PCB ban.^12^ Due to the wide variety of old and new products containing PCBs and the lack of natural degradation pathways for many PCBs, there continues to be widespread contamination from both Aroclor and non-Aroclor sources in buildings made with PCB-containing materials.

A particularly vulnerable population to PCB exposure is school-aged children, as many schools still in use today were built prior to the PCB-ban and newer buildings are likely to contain pigments and cabinetry finishes that contain non-Aroclor mixtures of PCBs. Because PCBs are semi-volatile, adolescents are exposed through inhalation of PCBs while at school. Studies of school air have found significantly elevated levels of PCBs in the air of many schools.^13,14^ One study found concentrations inside schools were 10-100 times higher than outdoors.^15^ While attending these schools, children accumulate PCBs. One study found that lower-chlorinated PCBs were detected in 95% of pupils attending a contaminated school compared to only 27% of the students at a non-contaminated school.^14^ Further rat studies using PCB mixtures similar to those found in schools (a combination of Aroclor 1254 and 1221) have reported PCB accumulation in tissues such as the liver and adipose tissue.^16,17^ Since children continue to be exposed to these PCB sources in their day to day lives, there is a dire need to understand how exposure to PCB mixtures affects lipid rich tissues, such as adipose.

While often thought of as just a storage depot for fat, adipose communicates with multiple other organ systems through endocrine signaling and plays a critical role in regulating whole-body energy metabolism. For example, adipose tissue stores excess lipids not only for caloric reserve, but also to avoid ectopic fat deposition in other organs such as the liver, muscle, and heart which would promote systemic complications such as non-alcoholic fatty liver disease, diabetes, and heart disease.^18^ Adipose tissue also releases adipokines, like adiponectin, which enhances insulin sensitivity and suppresses production of inflammatory cytokines such as TNFɑ.^19^ Thus, disruptions to adipose tissue metabolism and/or endocrine signaling can result in metabolic syndromes such as obesity, diabetes, and hyperlipidemia.^20^ During adolescence adipose tissue undergoes massive expansion via the proliferation and differentiation of adipose progenitors, also called adipose mesenchymal stem/stromal cells (MSC). Although adipose MSCs are responsible for maintaining healthy turnover rates of adipocytes throughout a person’s lifespan, these progenitor cells are particularly important during adolescence, when the number of adipose cells more than quadruples before becoming relatively constant in adulthood.^21^ Additionally, these adipose MSCs are active contributors to regulating local inflammation, particularly in the early stages of obesity.^22,23^ Early in the development of metabolic syndromes, adipose MSCs will produce high levels of MCP-1 leading to increased immune cell infiltration and inflammation.^24^ Other studies have shown stimulation of adipose MSCs is critical to regulating adipose inflammation.^25^ Since proper tissue expansion and immune function is imperative for adipose health, disruption of these adipose MSCs by environmental toxins would lead to disruption of metabolic health as the adipose becomes less able to replace adipocytes or control adipose inflammation.^20,26,27^ Therefore, it is imperative to understand if PCBs directly impact adipose MSCs. While many studies have been performed on adipocytes or pre-adipocytes exposed to PCBs, to date, the effects of PCB exposure on primary human adipose MSC has not been previously evaluated.^28^

Herein we systematically analyze how three different PCB mixtures impact human adipose MSCs. Two of the mixtures are found in legacy sources, Aroclor 1016 and Aroclor 1254, while the third mixture has been derived to mimic the PCB congeners recently found to be emitted from new cabinetry, Cabinet Mixture.^9,29^ All three mixtures were recently identified in a study looking at room-to-room variations in PCBs. PCB 47, the primary congener found in Cabinet Mixture, was identified in rooms built after 2012. Whereas rooms built before 1970 had evidence of Aroclor 1016 and 1254 likely from the use of fluorescent light fixtures and caulking respectively.^5,15^ Not only are these mixtures relevant to modern human exposure, but they also represent a wide range of congeners. Aroclor 1016 is comprised of primarily lower-chlorinated, non-dioxin-like PCBs (Dioxin TEQ: 0.09)^30^; Aroclor 1254 is comprised of higher-chlorinated PCBs including several dioxin-like congeners (Dioxin TEQ: 21)^30^; Cabinet Mixture is three non-dioxin-like congeners.^9,31^ A range of concentrations of each mixture are used to assess how short-term exposure impacts human adipose MSC growth, viability, and functional phenotype.

## METHODS

### Materials

#### Sources of PCBs

Aroclor 1016 and Aroclor 1254 (lot number KC 12-638) in the original containers from Monsanto (St. Louis, MO) were provided by the Synthesis Core of the Iowa Superfund Research Program (ISRP). The PCB congener profiles of both Aroclors have been reported previously.^32,33^ The cabinet PCB mixture was prepared by mixing 2,2’,4,4’-tetrachlorobiphenyl (PCB 47), 2,2’,4,6’-tetrachlorobiphenyl (PCB 51), and 2,3’,4,5’-tetrachlorobiphenyl (PCB 68) from AccuStandard (New Haven, CT, USA) in a weight ratio of 75:17:8. The original data and characterization of cabinet mixture are openly available through the Iowa Research Online repository at https://doi.org/10.25820/data.006184.

#### Cell culture media

Unless otherwise specified, cells were cultured in MEM-alpha (Thermo Fisher, Cat#: 12561049) supplemented with 1% (v/v) penicillin/streptomycin (Life Technologies), 1% (v/v) L-glutamine (Life Technologies), and 0.5% or 15% fetal bovine serum (VWR) depending on the experiment. For differentiation of adipose MSCs to adipocytes, two additional media formulations were used. Initiation of differentiation was done with Preadipocyte Differentiation Media (PDM-2) (Lonza, Cat: #PT-8002). Maintenance of differentiation was done with DMEM supplemented with 1.9 ng/mL Insulin (Sigma-Aldrich, Cat: #91077C) and 10% FBS.

### Isolation and characterization of adipose-derived MSC

MSCs were isolated from the stromal vascular fraction of human adipose. Briefly, adipose tissue from three breast reduction surgeries, donors 20-40 years of age, were obtained from the University of Iowa Tissue Procurement Core. The core collects tissue specimens from surgeries performed at the University of Iowa Hospitals and Clinics after obtaining informed consent according to an approved IRB held by the core. The core then removes any identifying information and provides the de-identified tissue to researchers. Once the tissue was obtained, adipose was dissected out, minced into small pieces, and incubated overnight in collagenase. The next day, the tissue was further disrupted via serial pipetting and centrifuged to separate the stromal vascular fraction from the lipid-rich layer. The SVF was collected, washed 3 times, and plated in polystyrene flasks with MEM-alpha growth media supplemented with 15% FBS. 4 hours after plating, any unattached cells were discarded, and the remaining cells were cultured until 70% confluent. The cells were then passaged 1:3 and expanded into a P1 generation for cryobanking and analysis of surface markers and differentiation potential.

To determine if the isolated cells were indeed MSCs, they were tested for conformance to the MSC minimal criteria.^34^ Cells between passage 1 and 2 were stained for CD90, CD73, CD105 CD34, CD45, CD11b, CD19, and HLA-DR surface expressions. Positive surface marker expression staining was carried out using PE-CD90 antibody (BD Biosciences, A15794), PE. Cy7-CD73 antibody (BD Biosciences, Cat #561258), and FITC-CD105 antibody (BD Biosciences, Cat #561443) with their corresponding isotype controls: PE-CD90 Mouse IgG1 (Invitrogen, Cat #GM4993), PE. Cy7 Mouse IgG1k (BD Biosciences, Cat #557872), and FITC Mouse IgG1k (BD Biosciences, Cat #56649) respectively. CD34, CD45, CD11b, CD19, and HLA-DR were assessed using a PE-conjugated hMSC Negative Cocktail (BD Biosciences, Cat #562530). After staining, cells from each donor were analyzed on a Cytek Northern Lights Spectral Cytometer **(Supplemental Figure 1)**.

**Figure 1:**
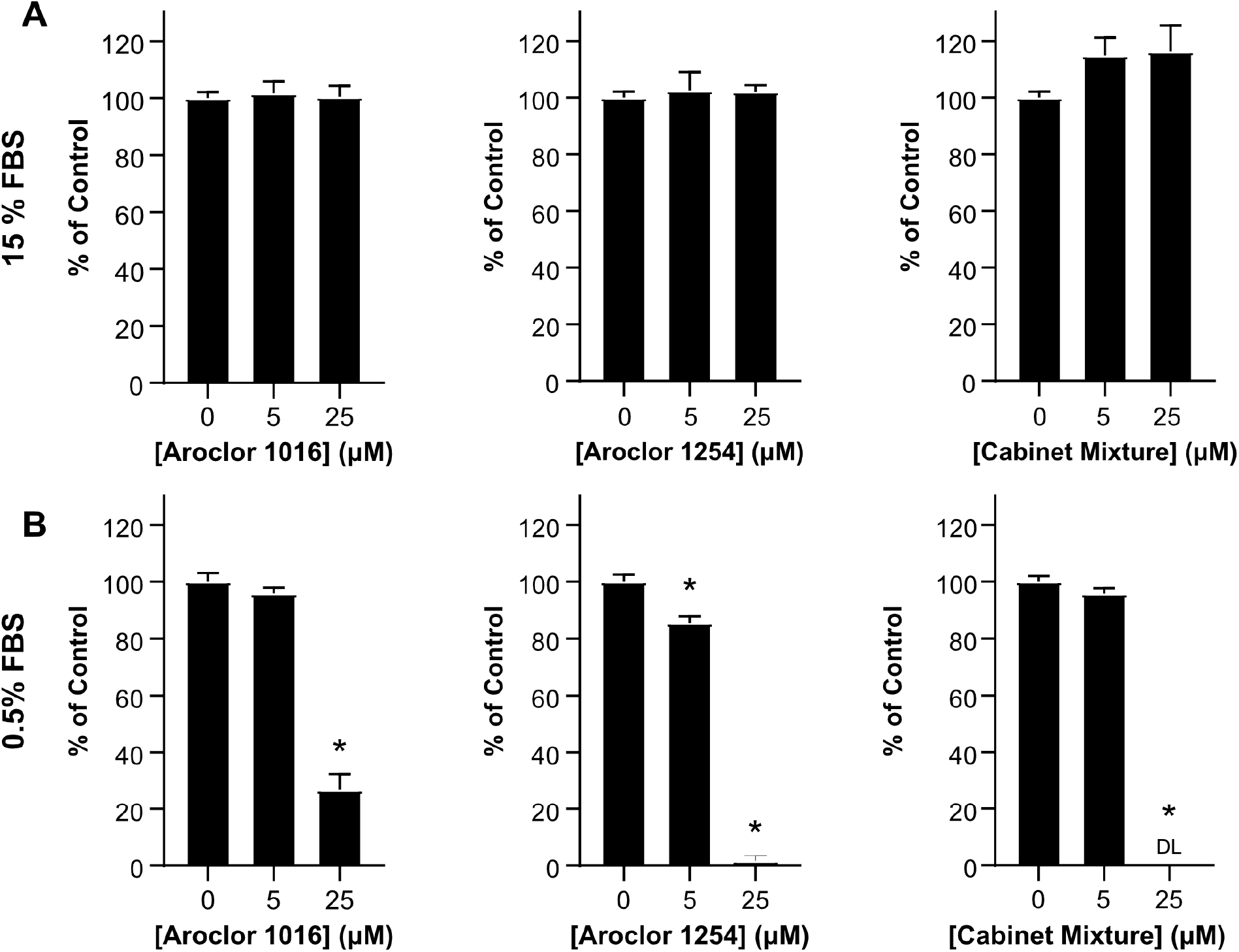
The effect of PCBs on MSC metabolic activity is highly dependent on serum concentration. The metabolic activity of MSCs was determined using XTT after 48 hours of exposure to 5 or 10 μM concentrations of Aroclor 1016, Aroclor 1254, or Cabinet Mixture dissolved in (A) 15% FBS media or (B) 0.5% FBS media. Bars represent mean and error bars are SEM. Ordinary one-way ANOVA, * designates significant difference (p<0.05) between the indicated group and vehicle control (0 μM) treated MSCs after Dunnett multiple comparison corrections. n=8 biological replicates with adipose MSC donor 2334 (representative of 4 experiments).

### Metabolic Function Assay

To determine the toxicity of the PCB mixtures on MSC metabolic function, we used an XTT assay (Biotium, Cat #30007). Using either 15% FBS or 0.5% FBS media, 2,000 MSCs were plated in 96-well plates containing 167 μL of media with a DMSO control (1 μL/mL) or media with 5 or 25 μM of PCBs dissolved in DMSO. After 48 hours, the culture media was removed and replaced with 100 μL of 15% FBS or 0.5% FBS media, depending on the original media composition. Then, 50 μL of XTT solution was added to the 100 μL of media in each well followed by incubation at 37 °C for 2 hours. After incubation, absorbance was read at 490 nm and 650 nm. Wells without MSCs and MSCs that had been permeabilized with 0.2% TritonX-100 were used as internal controls for each experiment. To show the change in XTT signal as a percent of the vehicle treated controls, all samples divided by the average of the vehicle treated controls.

### Cell Proliferation and Morphology

Cell proliferation was determined by counting nuclei stained with Hoechst 33342 (Thermo Fisher Scientific, Cat # H3570) and morphology was evaluated using Hoechst 33342 and ActinGreen 488 (Thermo Fisher Scientific, Cat # R37110) after 48 hours of PCB exposure. MSCs were seeded on 24-well plates with a seeding density of 12,000 cells/well. One mL of 0.5% FBS media with a DMSO control (1 μL/mL) or 0.5% FBS media with 1, 5, 10 20, 25 μM of PCBs dissolved in DMSO was added to each well at the same time as seeding. After 48 hours of incubation, media was removed from each well. The MSCs were then fixed for 5 minutes using 10% formalin followed by permeabilization with 0.05% Triton X-100 in PBS. A staining solution was made by diluting Hoechst 33342 and ActinGreen 488 with PBS to concentrations of 5 μL/mL and 2 drops/mL, respectively. After washing the cells with PBS, the staining solution was added to each well, and cells were incubated for 30 minutes at room temperature. The staining solution was removed and replaced with PBS. Imaging was performed at 10x magnification on an inverted fluorescent microscope (Leica DMI6000). To prevent bias in field selection, 5×5 tile scans were performed around the center of the well (25 images per well). The number of nuclei was counted using ImageJ (NIH) with an automated cell counting macro that ran a Gaussian blur, threshold, convert to mask, and watershed before analyzing particles to obtain the number and size of nuclei.

### Cell Viability

Cell death was assessed using propidium iodide staining and the LDH-Glo Cytotoxicity Assay (Promega, Cat #J2380). For the propidium iodide staining, MSCs were seeded at 61,000 cells/well in a 6-well plate. The MSCs were cultured for 48 hours in 5 mL of 0.5% FBS media with a DMSO control (1 μL/mL) or 0.5% FBS media with 1, 5, 10, 20, 25 μM of PCB mixtures dissolved in DMSO. The cells were then lifted and stained with propidium iodide (Sigma-Aldrich, Cat # P4864) to a final concentration of 0.01 μM. Controls included unstained MSCs as well as dead control cells which had been permeabilized with 0.2% TritonX-100 for 10 minutes. The cells were then incubated in the staining solution for 10 minutes before being analyzed via flow cytometry using a Cytek Northern Lights spectral cytometer. For the LDH assay, MSCs were plated in a 24-well plate as described in the “Cell Proliferation and Morphology” section. After 48 hours of incubation, 2 μL of media was collected from each well and each sample was diluted with 48 μL of LDH storage buffer. LDH Detection Reagent was prepared and added to each sample as directed by the manufacturer’s protocol. After incubation for 1 hour at room temperature, luminescence was recorded and normalized by the positive control to obtain LDH release as a % of the dead cell control.

### MSC-PBMC Direct Contact Co-culture

Peripheral blood mononuclear cells (PBMCs) were isolated from a leukapheresis reduction cone from a de-identified donor via the DeGowin Blood Center at the University of Iowa Hospital and Clinics. PBMCs were cryopreserved in a solution of 40% FBS, 50% RPMI, and 10% DMSO until use. Immunosuppressive capabilities of MSC were investigated utilizing a co-culture method as previously described.^35^ MSCs in a T75 flask were exposed to 1 μL/mL DMSO (“Vehicle Control”), 5, or 10 μM of PCB mixtures in 0.5% FBS containing media. Cells were then cultured within these conditions for another 48 hours. After pre-exposure to PCB mixtures, MSCs were harvested, counted, and plated for co-culture. PBMCs were stained with CFSE Cell Division Tracker dye (BioLegend; Cat: #423801) for 15-minutes. Any unbound CFSE dye was quenched with RPMI (15% FBS). Two-hundred fifty thousand cells were then added to each well to establish a 1:3 ratio of MSC to PBMCs. To activate PBMCs, 250,000 CD3/CD28 Dynabeads (Thermo Fisher Scientific, Cat: #11132D) were added to each well. A stimulated control with Dynabeads but no MSCs and an unstimulated control without MSCs or Dynabeads were performed in parallel and used for gating and statistical comparison. After 4-days of co-culture, PBMCs were collected and analyzed by flow cytometry. Gates were first set on FSC-A vs SSC-A for excluding any Dynabeads, MSCs, and debris within the samples. The resultant cells were then analyzed for percent proliferation as measured by CFSE intensity using the unstimulated control to set a gate.

### Adipogenic Differentiation Assay

To investigate the PCB mixtures’ potential disruption on adipogenesis through pre-exposure, cells were plated at a confluent density and cultured with 0.5% FBS MEM-alpha with 1 and 10μM of Aroclor 1016, 1254, or Cabinet Mixture for two days to achieve confluency. As a vehicle control, cells were cultured in 0.5% MEM-alpha containing 1μM of DMSO. After the pre-exposure period, all wells were washed with 1x PBS to remove remaining PCBs, and media was switched to 10% FBS Preadipocyte Differentiation Media (PDM-2) for 7 days, with media changes every 3-4 days. After 7 days the media was switched to 10% FBS DMEM + 1.9 ng/mL Insulin for another 7 days, with media changeouts every 3-4 days. As a negative control, cells received complete 0.5% FBS DMEM for all 14 days. At the end of the 14 days, media was collected for adiponectin analysis and cells were either stained with AdipoRed and imaged or harvested for RT-qPCR analysis.

#### Adiponectin ELISA

Media was collected after 14-days of differentiation and stored at −20 C until analysis. Quantification of adiponectin production was performed using an ELISA kit (BioLegend; Cat #442304) with no dilutions of samples to provide absorbance values within the linear range of the standard curve. Four biological replicates were used for each condition.

#### RT-qPCR Analysis

In preparation for RNA isolation, all samples were lifted with Accutase, spun down at 500g for 5 minutes, and washed with PBS. Total RNA was isolated via RNeasy kit (Qiagen; Cat #74104) as per the manufacturer’s protocols and eluted in 50 μL nuclease-free water. Each RNA elution was run on nano-drop for nucleotide quantification and ensuring protein/organic solvent purification. cDNA was synthesized utilizing a high-capacity cDNA reverse transcriptase kit (Applied Biosystems; Cat #4375575). ABI QuantStudio (model 7 Flex) was used for quantitative PCR reactions with SYBR green master mix (Applied Biosystems; Cat #4367659). The catalog of chosen primers can be found in supplementary data **(Supplementary Table 1)**. GAPDH was chosen for normalizing gene expression, and fold changes were compared to fully differentiated vehicle controls via 2^-ΔΔCt^ method.

### Statistical Analysis

GraphPad Prism 9 was used for graphing and performing statistical analyses on quantitative data. One-way ANOVA with Dunnett post-hoc analysis was used for statistical comparisons. A p<0.05 was considered statistically significant for post-hoc analyses. IC50 values were determined using Nonlinear Regression (least squares fit) for the [Inhibitor] vs. response (three parameters) model. Bottom constraint was set to 0 and top constraint was set to 152, the average number of cells/frame for the vehicle control condition. Further statistical details are provided within each figure caption.

## RESULTS AND DISCUSSION

### Toxicity of PCB mixtures is heavily influenced by serum

To investigate the effect of the PCB mixtures on MSC health, we decided to first determine if PCBs are toxic to MSCs. For assessing toxicity, MTT and XTT assays are often a first choice as they can detect a broad range of cellular responses. Since these assays measure NADH production, changes in the viability, metabolic function, or proliferative capacity of the cells will all lead to a change in signal. During our first XTT experiment, we performed a 48-hour exposure of MSCs to PCB mixtures dissolved in 15% FBS media. We found the PCB mixtures had little to no effect on the MSCs at both 5 and 25 μM concentrations **(Figure 1A)**. These data were surprising since previous work has shown various PCBs negatively impact many cell types such as human preadipocytes, neural stem cells, and astrocytes.^36–38^ Therefore, we were expecting to see a similar negative impact of the PCB mixtures on MSCs.

We were curious if there was a component of our cell culture system preventing PCBs from exerting an effect on MSCs. Previous work shows the drug binding sites of albumin strongly bind PCB congeners.^39–41^ For our first experiment, 15% of the media was FBS, of which almost half of the proteins were albumin.^42^ We hypothesized that the albumin in our media was significantly decreasing the amount of free PCB available to MSCs, thus masking any toxic effects of PCB exposure. To test this hypothesis, we repeated our first experiment, exposing the MSCs to the PCB mixtures for 48 hours, however, we replaced the 15% FBS media with a 0.5% FBS media.

After 48 hours of PCB exposure in this 0.5% FBS media, we found striking differences between the vehicle control and 5 and 25 μM conditions. Aroclor 1254 had close to a 20% reduction in NADH production at 5 μM, and all three PCB mixtures had 75% or greater reductions in NADH production at 25 μM exposures **(Figure 1B)**. Compared to the 15% FBS media, the 0.5% FBS media allowed for much greater insight into the effects of PCBs on MSCs and revealed PCB exposure disrupts adipose-derived MSCs cellular processes. Thus, all subsequent cell experiments were performed using MEM-alpha with 0.5% FBS unless otherwise stated.

### Short term exposure to PCB mixtures reduces adipose derived MSC expansion

While the XTT assay showed there was decreased NADH as the amount of PCB increased the decrease could have been due to direct PCB cytotoxicity, decreased proliferation, or decreased cellular metabolism. We next wanted to determine which of these potential mechanisms of cellular disruption were at work. Throughout the course of the XTT assay, we observed that the number of MSCs visible in the wells under a microscope at the end of the experiment was decreased in the PCB exposed conditions compared to the vehicle control. These observations of led us to hypothesize that increasing MSC exposure to PCBs would lead to decreased cell numbers. To determine the effect of PCBs on MSC expansion, we exposed MSCs from three donors to the three PCB mixtures at concentrations ranging from 1-25 μM. After 48 hours of incubation, we stained the cells with Hoechst and Actin Green and imaged them to assess cell counts and morphology.

We found that increasing concentrations of PCB mixtures led to a significant decrease in the number of cells in each well. All three MSC donors had significant decreases in cell counts with exposure to increasing concentrations of all three PCB mixtures **(Figure 2B)**. In fact, all samples exposed to 20 μM of PCBs had at least a 90% reduction in the number of MSCs at 48 hours.

**Figure 2:**
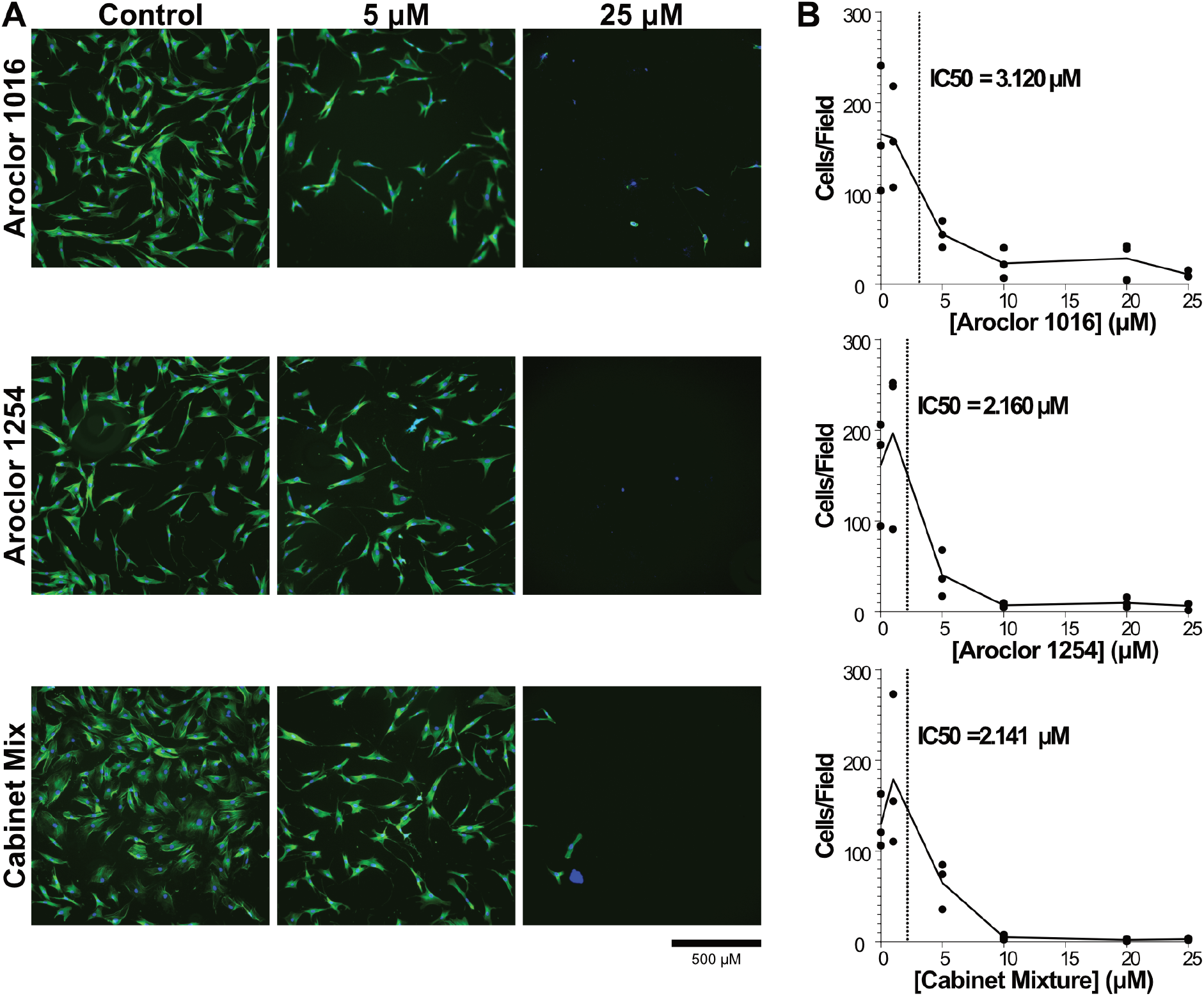
Cell count decreases with increasing exposure to PCB mixtures. (A) Representative images of adipose MSCs exposed to 0, 5, or 25 μM concentrations of Aroclor 1016, Aroclor 1254, and Cabinet Mixture. (B) The number of cells was determined by using ImageJ software to count the number of nuclei per image field. Quantification was performed on 25 images per condition for three independent adipose MSC donors. Data are represented via a point for the mean of each donor (n=25 image fields/point) with a line connecting the mean value of all donors (n=3 donors). IC50 values calculated using Prism for Aroclor 1016, Aroclor 1254, and Cabinet Mixture are 3.120 μM, 2.160 μM, and 2.141 μM, respectively.

Combining all donors together, IC50 values for each PCB mixture were calculated to be in the 2-5 μM range showing that even low concentrations of Aroclor 1016, Aroclor 1254, and Cabinet Mixture can significantly impact MSC health. In addition, there were also significant changes to the morphology of the MSCs with increasing concentrations of PCB mixtures. The cells that remained in the wells at the high concentrations (20, 25 μM) had what looked to be only fragments of cytoskeleton left and the portion that remained took on a spindle-like appearance and smaller overall footprint compared to cells in the vehicle control **(Figure 2A)**. In addition, the size of cell nuclei generally became larger and the variability of nuclear size increased **(Supplemental Figure 2)**. It should be noted that due to wash steps in the staining procedure, any small nuclei of detached dead cells would have been washed out before imaging. Of the cells that remained attached, the increased frequency of large nuclei at higher concentrations could be an indication that the cells have entered senescence, which is characterized by large, flattened nuclei.^43^

### PCB mixtures increase cell death at high concentrations

After the XTT and imaging studies, it was clear that exposure to PCB mixtures was causing cytotoxicity, but it was not yet clear if this was due primarily to suppression of cell proliferation or increases in cell death. To assess cell death directly, we used two complimentary assays, a propidium iodide (PI) stain to measure membrane permeability and lactate dehydrogenase (LDH) assay to measure release of LDH from dead cells. We again exposed MSCs to three PCB mixtures ranging from 1-25 μM and compared them to DMSO treated control cells. After 48 hours, the cells were stained with PI for analysis by flow cytometry and the media was assessed for LDH activity.

Based on the prior imaging experiments **(Figure 2)**, we expected to see increases in cell death starting at 5 μM, however, with both the PI staining **(Figure 3A)** and the LDH assay **(Figure 3B)**, we observed minimal cell death after exposure to 1, 5, or even 10 μM of the PCB mixtures. It was only at high exposure levels, 20 and 25 μM, that we observed significant levels of cell death. Thus, at higher concentrations >20 μM, the decrease in cell numbers is due to lethal cytotoxicity of the PCBs on MSCs. While the decreased cell counts we saw with 5 and 10 μM exposure but without increased levels of cell death suggests that low concentrations of PCB mixtures cause a cytostatic rather than a cytotoxic effect on adipose MSCs. These results indicate that PCB mixtures at lower concentrations are disrupting cellular processes involved in cell proliferation and raise the possibility that exposure to low concentrations of PCB mixtures alter other aspects of adipose MSC phenotype.

**Figure 3:**
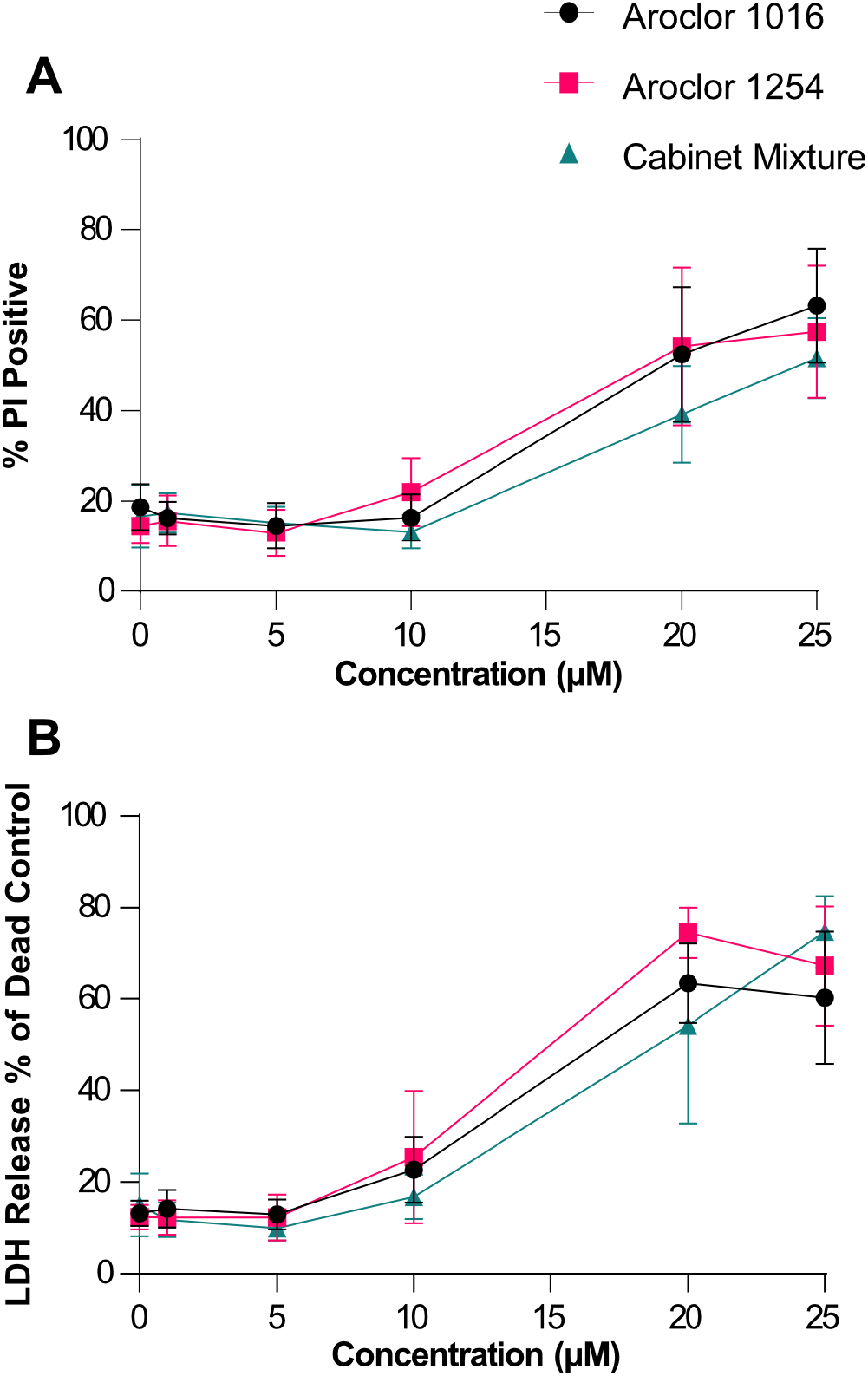
High concentrations of PCB mixtures kill the majority of adipose MSCs. Cell death was assessed using (A) propidium iodide staining and flow cytometry or (B) LDH assay after 48 hours of exposure to media formulated to contain 0, 1, 5, 10, 20, or 25 μM of Aroclor 1016, Aroclor 1254, and Cabinet Mixture. Each data point represents mean of 3 donors and error bars are SEM.

### Exposure to PCB mixtures only modestly impacts adipose MSC’s immunomodulatory properties

After determining the PCB mixtures’ cytotoxic effects on MSCs at higher concentrations, we wanted to investigate if exposure to non-lethal concentrations alters functional characteristics critical for adipose MSCs. An important property of adipose MSCs is their immunosuppressive capability. When in an inflammatory environment, MSCs tend to drive the surrounding immune cells towards a more immune-resolving phenotype, and as such serve as a key regulator of adipose inflammation.^25^ To determine how MSC exposure to non-lethal concentrations of PCB mixtures impacts their immunosuppressive properties, we pre-exposed the MSCs to each mixture at 5 or 10 μM for 48 hours within 0.5% FBS supplemented media. The cells were then washed to remove any dead cells and residual PCBs, counted, and seeded at a ratio of 1 MSC to 3 Dynabead-stimulated PBMCs (**Figure 4A)**.

**Figure 4:**
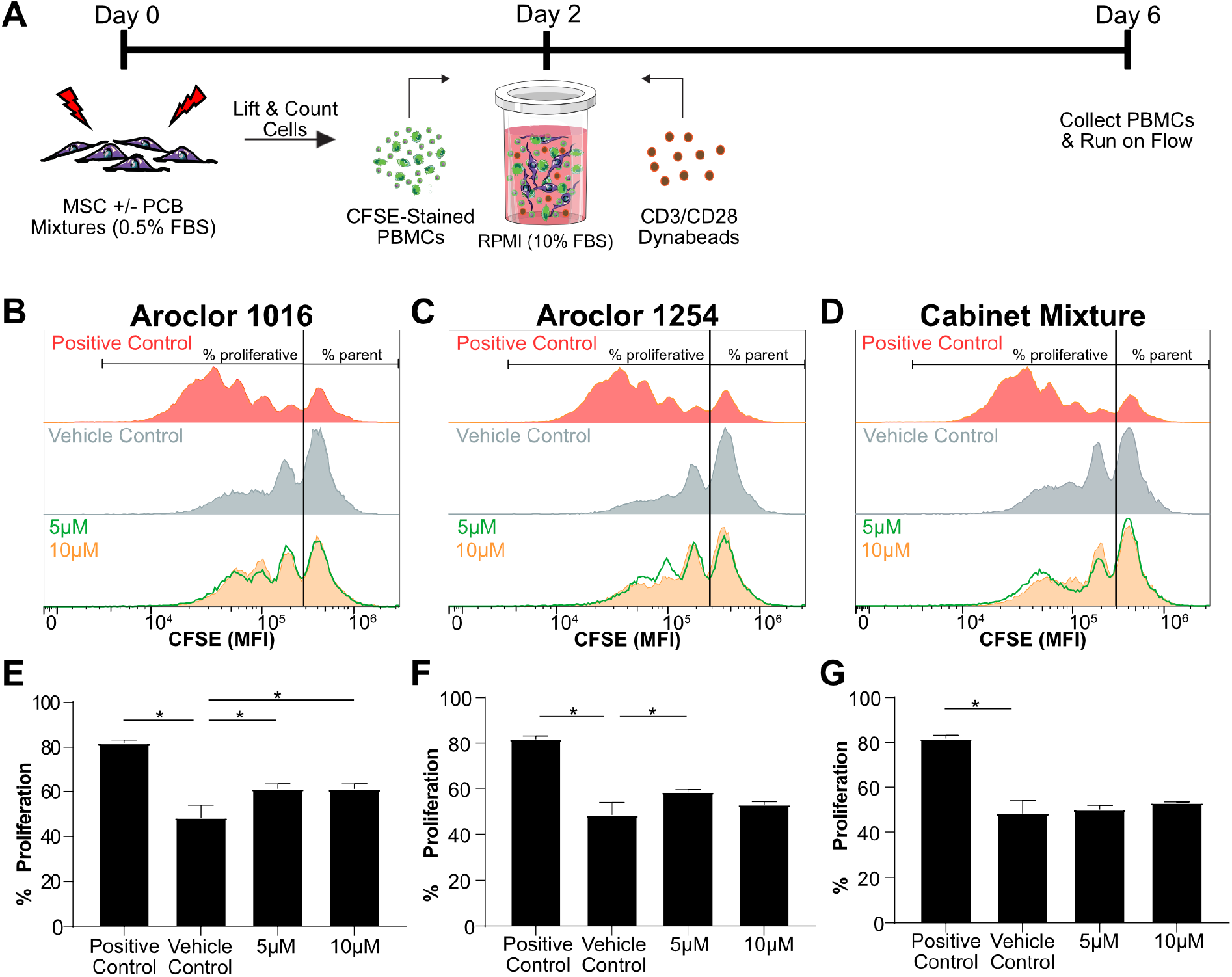
MSC immunosuppressive capabilities are only slightly reduced following exposure to PCB mixtures. A)Timeline of the PBMC-MSC coculture. Immunosuppressive capabilities of MSCs was assessed by measuring the dilution of CFSE dye in PBMCs after coculture with MSCs that had been pre-exposed for 48-hours to B) Aroclor 1016, C) Aroclor 1254 or D) Cabinet Mixture. The percent of PBMCs that proliferated was quantified to assess MSCs immunosuppressive potency after exposure to E) Aroclor 1016, F) Aroclor 1254 or G) Cabinet Mixture. Bars are mean +/-SD of n=3 independent adipose donors. Ordinary one-way ANOVA, * designates significant difference (p<0.05) between the indicated group and vehicle control (DMSO) pre-exposed MSCs after Dunnett multiple comparison corrections. Cell figures were adapted from https://smart.servier.com/ and licensed under CC-BY 3.0.

Since the PBMCs are stained with the cell proliferation dye, CFSE, prior to stimulation, each division will lead to a cytosolic partitioning of the fluorescent dye and leftward shift in CFSE intensity. As seen in the “Positive Control” panels of **(Figure 4B-D)**, the absence of MSCs allows the PBMCs to freely proliferate, leading to increased peaks at lower fluorescent intensities, each peak signifying a new generation of inflammatory cells. Upon adding vehicle treated MSCs, the divisions are substantially reduced. To summarize the degree of inflammatory suppression, we use a gate to separate the un-proliferated parent peak from cells that have undergone cell division and calculate the “% Proliferated”. The presence of vehicle control MSCs decreases the % of proliferated cells from ∼84% to ∼48% **(Figure 4E-G)**. Pre-exposure of MSCs to Aroclor 1016 at 5 and 10 μM both lead to a small but statistically significant increase in % proliferation (**Figure 4E)**. This was also observed for Aroclor 1254 **(Figure 4F)** at 5 μM, but the difference at 10 μM was not large enough to reach statistical significance. Interestingly, while both Aroclor 1016 and 1254 had modest effects on MSC suppression of PBMCs, Cabinet Mixture had no measurable effect **(Figure 4G)**. Based on prior cytotoxicity assays (**Figure 2**) that showed a change in cell behavior at these same concentrations, namely a dramatic reduction in proliferation, we expected to see a much larger impact on adipose MSCs immunosuppressive potency. This result paints a more complex portrait of adipose MSC response to PCB exposure and suggests the surviving adipose MSCs retain some functionality.

### Pre-Exposure to PCB Mixtures Disrupts MSCs’ Adipogenic Potential

With recent studies correlating persistent organic pollutants, such as PCBs, with the development of metabolic syndromes, we next wanted to investigate the influence of PCB mixtures on adipose MSCs adipogenic potential.^44^ To assess this, we pre-exposed MSCs to sublethal concentrations of PCB mixtures for 48 hours, and then induced adipogenic differentiation for 14-days in the absence of PCB exposure. After 14 days, we analyzed the transcript levels of key genes involved or indicative of adipocyte differentiation, namely, peroxisome proliferator activated receptor gamma (*PPARG*) a gene that plays a vital role as a master switch for the adipogenic differentiation pathway, contributing to protein transcription that influences lipid accumulation and insulin sensitivity, fatty acid binding protein (*FABP6*) a gene involved with fatty acid uptake and metabolism, which is a precursor to lipid production, and adiponectin (*ADIPOQ*) which is exclusively expressed by mature adipocytes for encoding adiponectin, a major protein involved in regulating whole-body metabolism.^45,46^ We found exposure to any PCB mixture led to a significant reduction in the transcript levels of both *PPARG* and *ADIPOQ* (**Figure 5A**). While significant, the magnitude of reduction of *PPARG* was fairly modest, with exposure leading to a 1.35-1.65-fold reduction compared to vehicle-treated controls. The reduction of *ADIPOQ* was more pronounced with 10 μM cabinet mixture exposure leading to a nearly 4-fold reduction in expression. Interestingly, no significant changes were observed for *FABP6*. Thus, exposure of adipose MSCs for just 48 hours has a long-term impact on gene expression, even after 14 days of differentiation in PCB-free media.

**Figure 5:**
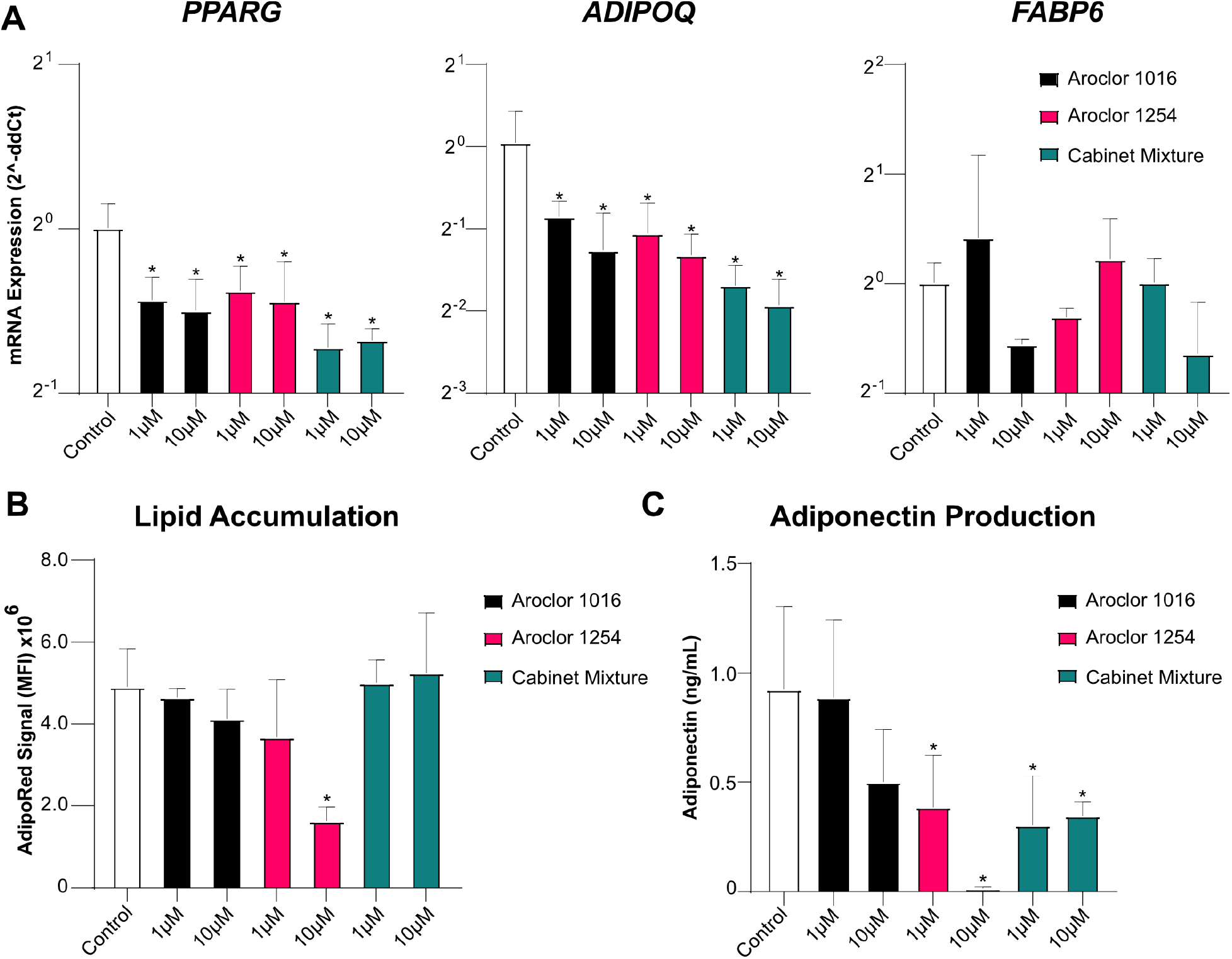
Adipogenic differentiation of MSCs are slightly diminished after pre-exposure to PCB mixtures. A) Fold-change of the expression of prominent genes in the adipogenesis signaling pathway, *PPARG, ADIPOQ*, and *FABP6*. Delta-CT was calculated using GAPDH and then compared to vehicle treated controls using the Delta-Delta-CT method. n=3 experiments with Adipose MSC donor 2334 B) Lipid accumulation as measured by AdipoRed staining measured using a 96-well plate reader. C) adiponectin present in culture media measured using ELISA. Bars are mean +/-SD of n=4 experiments with adipose MSC donor 2334. Ordinary one-way ANOVA with Dunnett post-hoc test, * indicates p<0.05 compared vehicle controls.

To determine if alterations in gene expression led to changes in adipocyte phenotype, we repeated the experiment and analyzed lipid accumulation and adiponectin production directly. MSCs exposed to Aroclor 1016 or 1254 showed a dose-dependent decrease in lipid accumulation while MSCs exposed to Cabinet Mixture showed no decline in lipid accumulation (**Figure 5B**). Interestingly, the only exposure that resulted in a statistically significant decline in lipid accumulation was 10 μM Aroclor 1254. Analysis of adiponectin secreted by the adipocytes after 14 days of differentiation revealed a stark decrease in production after pre-exposure to all three of the PCB mixtures (**Figure 5C**). Specifically, pre-exposure to Aroclor 1254 at 5 μM or Cabinet Mixture at both concentrations reduced adiponectin output in half. Increasing the Aroclor 1254 exposure to 10 μM nearly completely blocked adiponectin production. Overall, the observed changes in adiponectin secretion were consistent with the *ADIPOQ* transcript levels (**Figure 5A**). This is important, as adiponectin production assists in maintaining insulin-sensitivity and attenuating chronic inflammation, while deceased serum levels have been associated with obesity and type-2 diabetes.^47,48^ Taken collectively, these data demonstrate that even a short window of exposure to PCB mixtures disrupts the quality of adipogenesis which alters the properties of the resultant mature adipocytes.

## IMPLICATIONS FOR HUMAN HEALTH

Our work shows that exposure to Aroclor 1016, Aroclor 1254, and the Cabinet Mixture all result in adverse effects on adipose MSCs, a cell type that is critical for the maintenance and function of adipose tissue and overall health. We examined PCB concentrations that are similar to tissue concentrations measured in adipose. In vivo studies usually report PCBs in terms of ng/g of lipid and have found adipose tissue levels ranging from 700-9000 ng/g of lipid depending on the severity of exposure^49,50^. Considering the molecular weight of PCBs, the density of lipids, and that adipose is ∼60% lipids^51^, these tissue levels correspond to an adipose tissue concentration range of 1.5-18 µM of total PCB. While we examined mixtures rather than single congeners in this study, mixtures are more physiologically relevant as people are exposed to mixtures and not single congeners. Furthermore, studying mixtures made our assays sensitive to possible interaction effects between congeners which have been reported for other cell types^52^. Our data show that exposure to these mixtures at adipose tissue relevant concentrations results in significant toxicity and functional disruption of adipose MSCs, an important stem/mesenchymal cell population. In reality, PCB profiles found within current U.S. school air show contributions from all three of these investigated sources, but only specific congeners are detectable within the serum of students.^53^ Regardless, these findings have potential implications for the health of school-aged children, especially those attending schools that were built during Aroclor production. Adipose MSCs are particularly important during adolescence, as the number of adipocytes goes through massive expansion before plateauing and maintaining numbers throughout adulthood. This expansion of adipocytes relies on the MSC niche found within adipose tissue. We have shown here that exposure to non-lethal concentrations of PCB mixtures disrupts both adipose MSC proliferation and impairs their ability to differentiate into mature functional adipocytes. These effects of PCBs could have profound implications on the number and quality of adipocytes that are generated during expansion. Mature adipocytes with altered adiponectin signaling could have significant physiological effects such as decreased insulin sensitivity in peripheral tissues, disrupted androgen signaling, and increased chronic inflammation: all of which are aspects of metabolic syndrome.

While, in adults, MSCs and preadipocytes do not proliferate at the same rate as they do in adolescents, about 10% of all adipocytes are still replaced annually, and a healthy MSC niche is needed to support this replacement.^21^ Since PCBs are known to accumulate in adipose tissue, adipose MSCs within adults are susceptible to the negative effects of PCB exposure.^54,55^ Disruption of adipose MSCs would lead to a lower rate of adipocyte replacement which has been linked to hypertrophic obesity.^56^ Additionally, recent work suggests that proper expansion of adipose tissue is fundamental to the prevention of metabolic disease.^20,27^

While the effects of PCB mixtures on adipose MSCs were fairly consistent between different mixtures, there were also distinct differences between the groups. Exposure of MSCs to Aroclor 1254 leads to much lower levels of adiponectin and lipid accumulation compared to Aroclor 1016 or Cabinet Mixture. One likely explanation for this difference is the congeners which compose the mixtures. Aroclor 1254 contains both higher chlorinated and dioxin-like PCB congeners, while Aroclor 1016 contains lower chlorinated non-dioxin-like congeners.^32,33^ Moreover, each congener has its own partitioning coefficient and effective free concentration. Due to the differences in congener profile and relative abundance, it is likely that multiple distinct congeners, or congener subsets, are responsible for the biological effects we have observed here on adipose MSCs

Another potential reason for the differences between mixtures observed in the adipogenic differentiation assay is the different mechanisms of action of different PCBs. Previous work has focused primarily on elucidating the effect of dioxin-like PCBs on adipocytes. PCB 126, a dioxin-like PCB, activates the aryl hydrocarbon receptor (AhR) which suppresses PPARG transcription and, subsequently, adipogenesis.^36^ Further, transcriptome sequencing of adipocytes after exposure to PCB 126 showed marked activation of AhR genes but also activation of proinflammatory pathways and the AGE-RAGE pathways which is known to be associated with the development of obesity and insulin resistance.^57^ In our experiments, Aroclor 1254 is the only mixture that contains significant amounts of dioxin-like PCBs (Aroclor 1016 contains trace amounts of DL congeners but has a Dioxin TEQ of 0.09 compared to a TEQ of 21 for Aroclor 1254)^30^. Aroclor 1254 has also been shown to lead to higher levels of DNA damage, tumorigenesis, and disruption of central nervous system neurochemical function than Aroclor 1016.^3,58,59^ This disruption may be due to a dose-dependent inhibition of creatine kinase activity as seen in L6 myoblasts.^60^ Creatine kinase-B activity is decreased in adipocytes during obesity, thus opening the potential for a mechanistic tie between Aroclor 1254 exposure and adipocyte disfunction.^61^ The other two mixtures are comprised of non-dioxin-like congeners, thus the previous proposed mechanisms are unlikely to explain the effects of Aroclor 1016 and Cabinet Mixture. Non-dioxin-like PCBs, such as those found in Aroclor 1016 and Cabinet Mixture, have been shown to induce neuronal apoptosis via a p53-independent mechanism.^62^ They also bind to ryanodine receptors (RyR) and increase calcium release from the endoplasmic reticulum of neurons. Increased RyR activity is associated with neuronal apoptosis and dendritic growth.^62^ In adipocytes, RyR3 activity is inversely related to adiponectin mRNA expression, thus overactivation of RyR by non-dioxin-like PCBs is one potential mechanism of disrupted adipose function.^63^ While further studies will be required to understand which congeners or congener interactions are responsible for toxicity and their mechanisms of action, this work provides direct evidence that short-term exposure to environmentally relevant mixtures of PCBs disrupt the health and function of adipose MSCs.

**Supplementary Figure 1:**
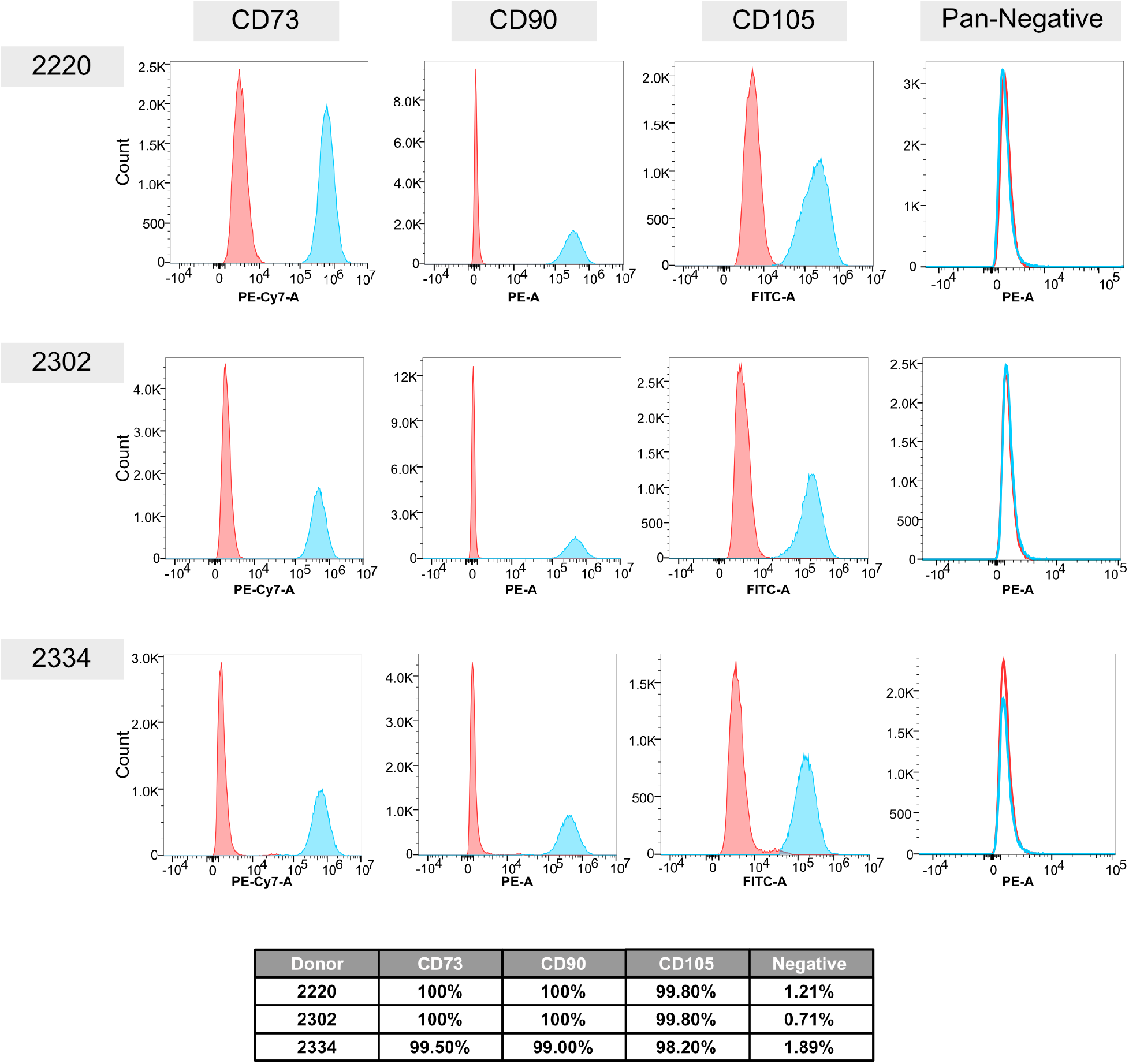
Adipose MSCs meet the ISCT Minimal Criteria. Adipose MSCs were analyzed for expression of markers according to the ISCT MSC Minimal Criteria. Flow cytometry histogram plots are shown for CD73, CD90, CD105 and a Negative Cocktail (CD34, CD45, CD11b, CD19, and HLA-DR). Each plot contains the isotype control (red peak) and on target sample (blue peak). Table displays the percent of each on-target population considered positive based on gates set using the isotype control.

**Supplemental Figure 2:**
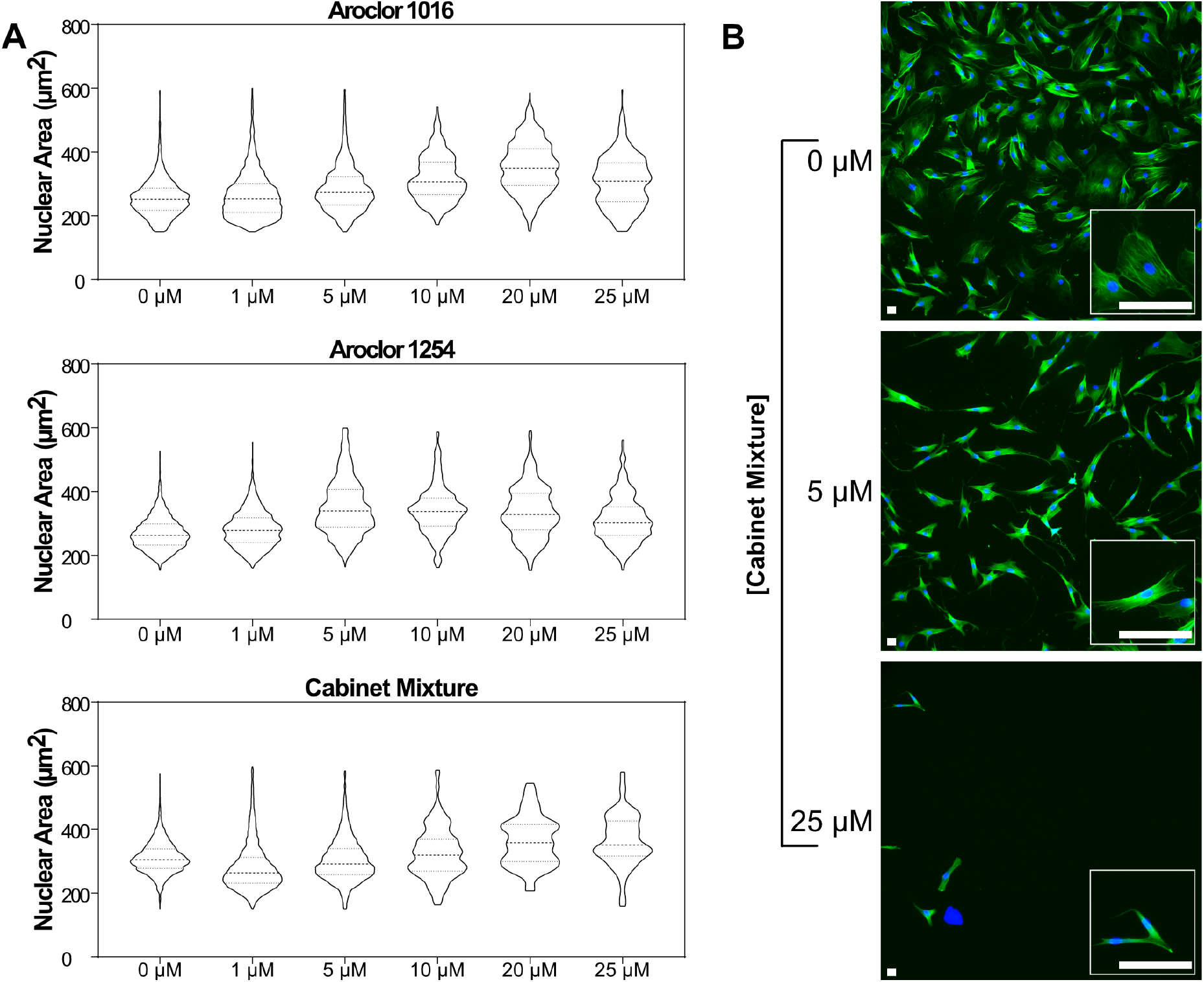
Exposure to PCB mixtures leads to dose dependent changes to MSC morphology. Morphology of nuclei and cytoskeleton after 48 hours of exposure to PCB mixtures. (A) Violin plot of nuclear area after exposure to 0, 1, 5, 10, 20, or 25 μM concentrations of Aroclor 1016, Aroclor 1254, and Cabinet Mixture. (B) Representative images of cells after exposure to 0, 5, or 25 μM concentrations of Cabinet Mixture. All scale bars represent 50 μm.

**Supplemental Table 1.**
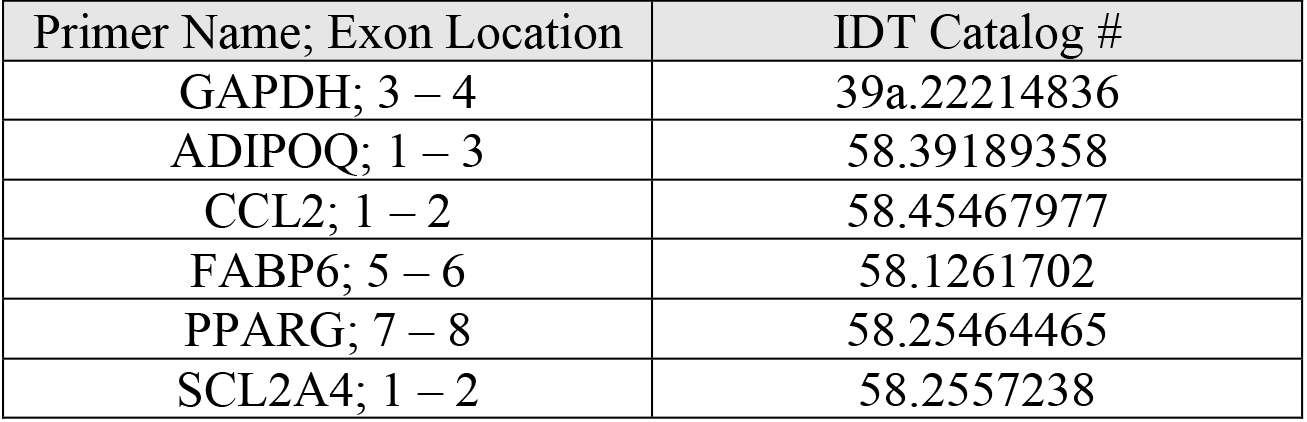
qRT-PCR Primers.

## ACKNOWLEDGEMENTS

This study was supported by NIH P42 ES013661 (JAA, AJK, HJL) and a pilot grant from the Environmental Health Research Center <u>P30 ES005605</u> (AJK, JAA). RB is supported by the University of Iowa MSTP Grant, NIH T32 GM139776. We also acknowledge the University of Iowa Tissue Procurement Core which manages the University of Iowa Biobank (UIBB – IRB#201103721) for services provided related to acquisition of study specimens. The TPC is supported by an award from NIH (NCI award number P30CA086862) and the University of Iowa Carver College of Medicine.

## AUTHOR CONTRIBUTIONS

Riley Behan-Bush: Conceptualization, Methodology, Formal analysis, Investigation, Data Curation, Writing – Original Draft, Writing – Review & Editing, Visualization. Jesse Liszewski: Conceptualization, Methodology, Formal analysis, Investigation, Data Curation, Writing – Original Draft, Writing – Review & Editing, Visualization. Michael Schrodt: Methodology, Investigation, Writing – Original Draft. Bhavya Vats: Investigation. Xueshu Li: Resources, Data Curation, Writing – Original Draft. Hans-Joachim Lehmler: Resources, Data Curation, Writing – Original Draft, Funding Acquisition. Aloysius J. Klingelhutz: Conceptualization, Resources, Writing – Review & Editing, Funding Acquisition. James Ankrum: Conceptualization, Methodology, Formal analysis, Resources, Data Curation, Writing – Review & Editing, Visualization, Supervision, Funding Acquisition.

