## Supplemental for "Toxicity impacts on human adipose MSCs acutely exposed to Aroclor and non-Aroclor mixtures of PCBs"

**Title: Exposure to Aroclor 1016, Aroclor 1254 and modern Cabinet Mixtures of polychlorinated biphenyls negatively impacts the growth, viability, and function of adipose stem cells.**

**Authors**
Riley M. Behan-Bush[1], Jesse N. Liszewski[1], Michael V. Schrodt, Bhavya Vats, Xueshu Li, Hans-Joachim Lehmler, Aloysius J. Klingelhutz, and James A. Ankrum[*]
[1]Contributed equally

**Supplemental Information**

**Supplementary Figure 1: Adipose MSCs meet the ISCT Minimal Criteria.**

**Supplemental Figure 2: Exposure to PCB mixtures leads to dose dependent changes to MSC morphology.**

**Supplemental Table 1. qRT-PCR Primers**

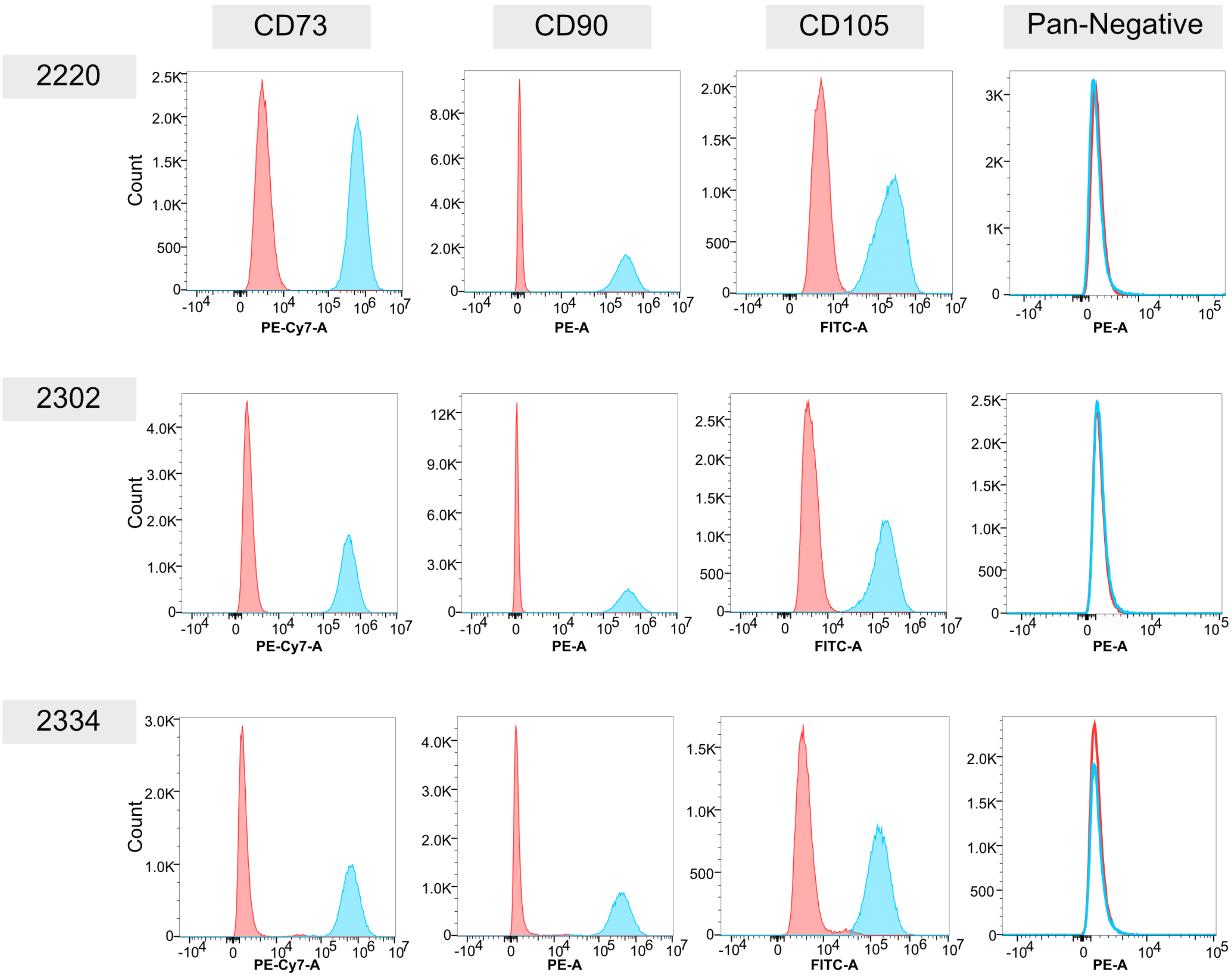


| Donor | CD73 | CD90 | CD105 | Negative |
|---|---|---|---|---|
| 2220 | 100% | 100% | 99.80% | 1.21% |
| 2302 | 100% | 100% | 99.80% | 0.71% |
| 2334 | 99.50% | 99.00% | 98.20% | 1.89% |

**Supplementary Figure 1: Adipose MSCs meet the ISCT Minimal Criteria.** Adipose MSCs were analyzed for expression of markers according to the ISCT MSC Minimal Criteria. Flow cytometry histogram plots are shown for CD73, CD90, CD105 and a Negative Cocktail (CD34, CD45, CD11b, CD19, and HLA-DR). Each plot contains the isotype control (red peak) and on target sample (blue peak). Table displays the percent of each on-target population considered positive based on gates set using the isotype control.

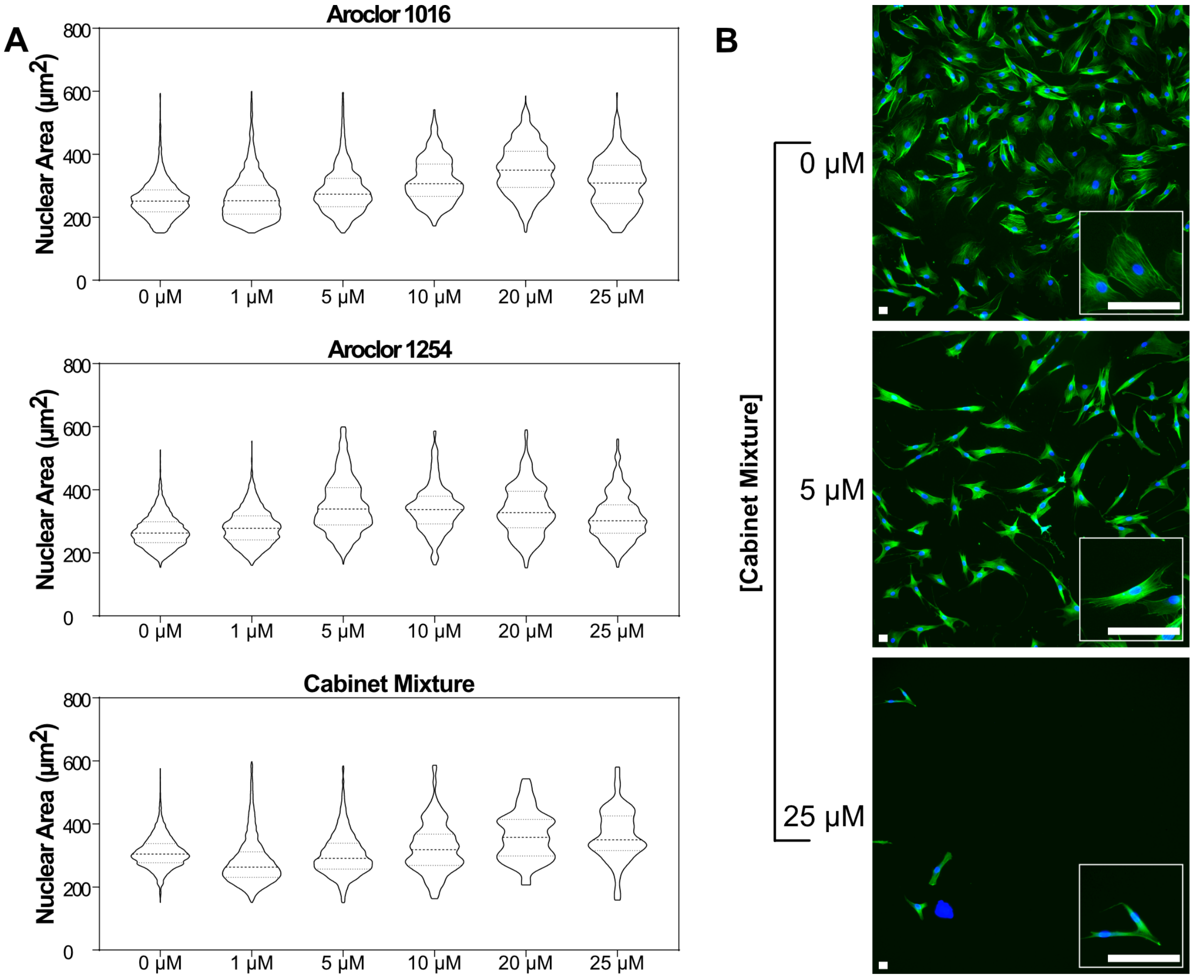


**Supplemental Figure 2: Exposure to PCB mixtures leads to dose dependent changes to MSC morphology.** Morphology of nuclei and cytoskeleton after 48 hours of exposure to PCB mixtures. (A) Violin plot of nuclear area after exposure to 0, 1, 5, 10, 20, or 25 µM concentrations of Aroclor 1016, Aroclor 1254, and Cabinet Mixture. (B) Representative images of cells after exposure to 0, 5, or 25 µM concentrations of Cabinet Mixture. All scale bars represent 50 µm.

**Supplemental Table 1. qRT-PCR Primers**

| Primer Name; Exon Location | IDT Catalog # |
|---|---|
| GAPDH; 3 – 4 | 39a.22214836 |
| ADIPOQ; 1 – 3 | 58.39189358 |
| CCL2; 1 – 2 | 58.45467977 |
| FABP6; 5 – 6 | 58.1261702 |
| PPARG; 7 – 8 | 58.25464465 |
| SCL2A4; 1 – 2 | 58.2557238 |
